# RAmbler: *de novo* genome assembly of complex repetitive regions

**DOI:** 10.1101/2023.05.26.542525

**Authors:** Sakshar Chakravarty, Glennis Logsdon, Stefano Lonardi

## Abstract

Complex repetitive regions (also called segmental duplications) in eukaryotic genomes often contain essential functional and regulatory information. Despite remarkable algorithmic progress in genome assembly in the last twenty years, modern *de novo* assemblers still struggle to accurately reconstruct these highly repetitive regions. When sequenced reads will be long enough to span all repetitive regions, the problem will be solved trivially. However, even the third generation of sequencing technologies on the market cannot yet produce reads that are sufficiently long (and accurate) to span every repetitive region in large eukaryotic genomes.

In this work, we introduce a novel algorithm called RAmbler to resolve complex repetitive regions based on high-quality long reads (i.e., PacBio HiFi). We first identify repetitive regions by mapping the HiFi reads to the draft genome assembly and by detecting unusually high mapping coverage. Then, (i) we compute the *k*-mers that are expected to occur only once in the genome (i.e., single copy *k*-mers, which we call *unikmers*), (ii) we barcode the HiFi reads based on the presence and the location of their uni*k*mers, (iii) we compute an overlap graph solely based on shared barcodes, (iv) we reconstruct the sequence of the repetitive region by traversing the overlap graph.

We present an extensive set of experiments comparing the performance of RAmbler against Hifiasm, HiCANU and Verkko on synthetic HiFi reads generated over a wide range of repeat lengths, number of repeats, heterozygosity rates and depth of sequencing (over 140 data sets). Our experimental results indicate that RAmbler outperforms Hifiasm, HiCANU and Verkko on the large majority of the inputs. We also show that RAmbler can resolve several long tandem repeats in *Arabidopsis thaliana* using real HiFi reads.

The code for RAmbler is available at https://github.com/sakshar/rambler.

**CCS CONCEPTS:** **Applied computing** → **Bioinformatics**; **Computational genomics**; *Molecular sequence analysis*; • **Theory of computation** → Graph algorithms analysis.

## 1 INTRODUCTION

Given the broad biological impact of obtaining the genome for a new organism, *de novo* genome assembly is one of the most critical problems in computational biology. Despite tremendous algorithmic progress, the problem is not yet completely “solved”. The assembly problem remains challenging due to the high repetitive content of eukaryotic genomes, short read length, uneven sequencing coverage, non-uniform sequencing errors and chimeric reads. Repetitive regions (or segmental duplications) are the primary reason *de novo* genome assemblies are often fragmented and incomplete. Large eukaryotic genomes often contain hundreds of long tandem repeats. For example, Figure 1 shows a frequency histograms of the repeats present in seven important plant species.

**Figure 1:**
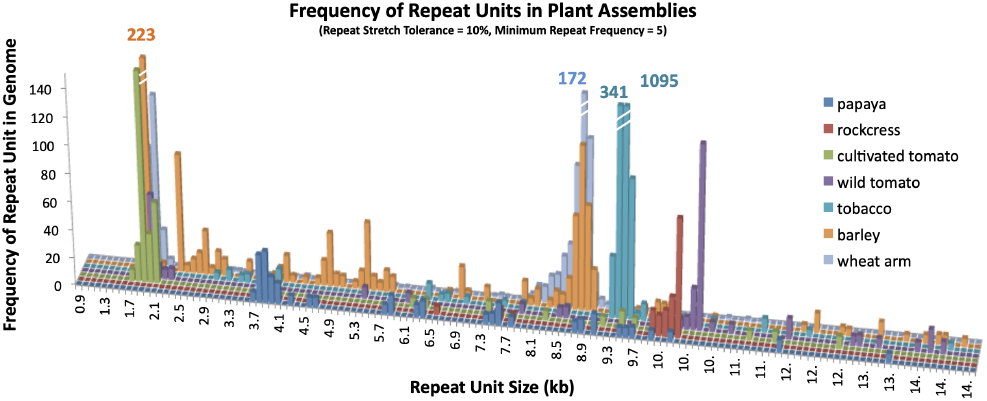
A frequency histogram of unresolved repeats in seven plant genomes (2015); the x-axis represents the repeat size (in kb); the y-axis represents the number of copies in each genome of those repeats’ sizes (figure reproduced with permission [3])

The third generation of sequencing technology on the market, e.g., Pacific Biosciences [5, 10, 12, 26, 31] and Oxford Nanopore [1, 4, 17, 27], offers longer reads at a higher cost per base than the second generation, but the sequencing error rate is much higher. The introduction at the end of 2019 of PacBio HiFi sequencing has been a “game-changer” in genome assembly, because it can produce read lengths averaging 10-25 kb with accuracy greater than 99.99% [37]. HiFi sequencing greatly improved human assemblies [6, 22, 25, 32]. The telomere-to-telomere human genome sequencing project took advantage of PacBio HiFi and Oxford Nanopore reads to close most of the repetitive gaps, and achieving 99.9% completeness [9, 16, 19, 21, 29]. In particular, the method developed to assemble human chromosome 8 critically depended on single copy *k*-mers to resolve repetitive regions.

In this work, (1) we carry out the first statistical analysis on the method to extract single copy *k*-mers for PacBio HiFi and Oxford Nanopore reads; (2) we develop a new assembly method, called RAmbler, that takes advantage of single copy *k*-mers to resolve complex repetitive regions; (3) we compare RAmbler against state-of-the-art assemblers (Hifiasm, HiCANU and Verkko) on more than 140 synthetic data sets; (4) we show that RAmbler can resolve complex repeats in *A. thaliana* from real HiFi data; (5) we introduce a new quality metric for genome assemblies that incorporates completeness, contiguity and number of mis-assemblies in one consolidated measure.

Due to its importance, the problem of reconstructing repetitive regions (segmental duplications) has been addressed several times in the literature, most recently in [2, 33]. However, the Segmental Duplication Assembler (SDA) proposed in [33] “*is no longer maintained and should not be used. SDA was designed for low quality PacBio (CLR) and ONT long reads, both of which are now substantially higher quality allowing standard assembly tools like Flye, HiCanu, and Hifiasm to outperform any results previously possible with SDA*.” (from the github page https://github.com/mrvollger/SDA).

## 2 METHODS

### 2.1 Problem Formulation

We assume that (i) the genome *G* contains *n* repetitive regions {*R*_1_, …, *R*_*i*_, …, *R*_*n*_}, (ii) each repetitive region *R*_*i*_ is composed of *t*_*i*_ tandem copies of a string *α*_*i*_, (iii) each tandem copy has sufficient variations that allows it to be distinguished from another copy; we assume that each copy contains SNPs with probability *p*_*SNP*_ (e.g., *p*_*SN P*_ = 1/ 100), and its length can increase or decrease by at most *L*%. Given a set *T* of HiFi reads and the draft genome *G*, the objective is to produce a set {*F*_1_, …, *F*_*i*_, …, *F*_*n*_} of *n* assemblies, where each *F*_*i*_ is as “similar as possible” to *R*_*i*_ . In particular, if the assembly *F*_*i*_ contains 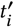 copies of the repeat unit, we want 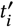as close as *t*_*i*_ as possible (see Figure 9). For synthetic data, we measure the quality of the assembly *F*_*i*_ by comparing it to *R*_*i*_ using QUAST [7], i.e., we report the fraction of *R*_*i*_ covered by *F*_*i*_ (ideally 100%), the number of mis-assemblies in *F*_*i*_ (ideally zero), and the number of contigs in *F*_*i*_ (ideally one). When the ground truth is unavailable (i.e., for real data sets), a qualitative assessment of the assembly’s accuracy can be obtained by aligning *F*_*i*_ with the corresponding repeat unit *α*_*i*_ . The alignment, visualized as a dot plot, can provide a qualitative measure on how well the repeat units are assembled.

### 2.2 Repeat Identification

To identify the repetitive regions {*R*_1_, …, *R*_*n*_}, we map the HiFi reads *T* against the draft genome assembly *G*. Since unresolved tandem repeats are collapsed in the draft assembly, they can be identified by a spike in mapping coverage. For instance, Figure 2 shows the mapping coverage of an unresolved tandem repeat in chromosome XII of *Saccharomyces cerevisiae* which is known to contain ∼150 tandemly repeated copies of a 9.1 kb rDNA unit [11, 13]. This region is the only unresolved non-telomeric gap in the current *S. cerevisiae* assembly.

**Figure 2:**
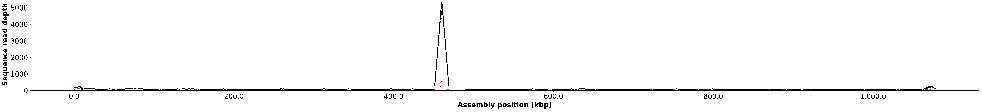
PacBio HiFi mapping coverage for chromosome XII in *Saccharomyces cerevisiae* illustrated using NucFreq; the coverage spike indicates the presence of a repetitive region which is known to contain ∼150 tandemly repeated copies of a 9.1 kb rDNA unit

### 2.3 Analysis of *k*-mer Distribution

Recall that we assume that the copies in the repetitive region are not identical to each other. If all the copies were identical, the problem of resolving repeats would be impossible, unless one can produce reads so long that they span the entire repetitive region. We rely on the presence of the SNPs to distinguish and partition the HiFi reads that belong to different repeat copies within a repetitive region. Since SNPs are uniformly distributed among the copies, we expect each copy to have its own “signature” of SNPs. Each SNP is likely to induce a unique (or single copy) *k*-mer, i.e., **a** *k***-mer that occurs a number of times approximately equal to the expected sequencing coverage**. We call these *k*-mers, *unikmers*. Uni*k*mers (called SUNK in [16]) were crucial to resolve the assembly of human chromosome 8, but to the best of our knowledge they have not been used in any other assembly method. Please note that uni*k*mers are NOT *k*-mers that appear only once in the reads: those *k*-mers correspond to sequencing errors.

The first contribution of our study is to provide a method to identify uni*k*mers from the reads, and analyze its accuracy and precision. RAmbler finds uni*k*mers by selecting all *k*-mers in the HiFi reads that have a number of occurrences within the interval [*μ*− *tσ, μ* + *tσ*], where *μ* is the average sequencing depth of the HiFi reads, *σ* is the standard deviation of the sequencing depth, and *k* and *t* are user-defined parameters.

We investigate how to choose *k* and *t* in the following analysis. (1) We determine the set of true uni*k*mers in the *Saccharomyces cerevisiae* genome to serve as the ground truth. (2) We compute the *k*-mer distribution for a set of real HiFi reads (SRA accession SRR 13577847) and Oxford Nanopore (ONT) reads (SRA accession SRR 15597407) for *S. cerevisiae* using Jellyfish [18]. Figure 3 shows the *k*-mer distribution for HiFi reads for *k* = 17 (left), *k* = 21 (middle) and *k* = 25 (right). Odd integers in the range [17, 25] are typical choices for *k* to estimate genome size (see, e.g., [28, 35]) or the construction of the de Bruijn graphs for eukaryotic genomes (see, e.g., [38]). Observe that the distributions are almost identical, which indicate that any of these *k*-mer choices would be appropriate. We compute the average sequencing depth *μ* and the standard deviation of the sequencing depth *σ* from the *k*-mer distribution. The *k*-mers in the HiFi reads that have a number of occurrences within the interval [*μ* −*tσ, μ* + *tσ*] for *t* = 0, 1, 2, 3, 4, 5 are compared against the true uni*k*mers: true positive, false positive, true negative and false negative are recorded.

**Figure 3:**
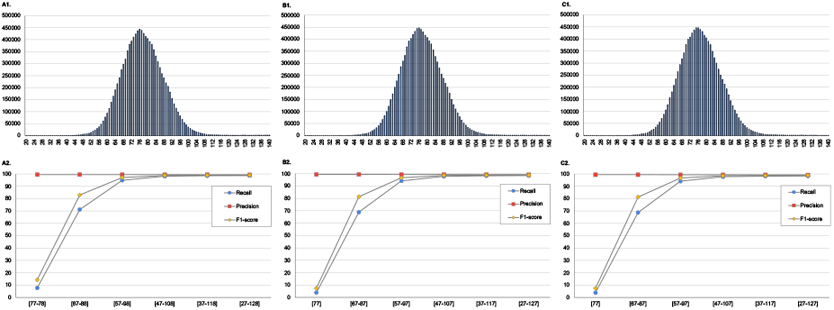
PacBio HiFi *k*-mer distribution for *k* = 17 (A1), *k* = 21 (B1) and *k* = 25 (C1) for *Saccharomyces cerevisiae*; precision, recall, and F1-score for uni*k*mers when *k* = 17 (A2), *k* = 21 (B2) and *k* = 25 (C2) for larger and larger intervals centered at the average sequencing coverage

The results of this analysis (precision, recall and F1-score) for HiFi reads are show in the bottom row of Figure 3. The x-axis represents the choice of *t*, i.e., larger and larger intervals centered around the mean (the first interval is for *t* = 0, the second is for *t* = 1, etc.). Figure 3-A2 shows the results for *k* = 17, Figure 3-B2 illustrates the results for *k* = 21 and Figure 3-C2 shows the results for *k* = 25. Observe that in all cases precision and recall are very close to 100% as soon as *t* = 3. For instance, there are 11,137,337 21-mers that occur [47 − 107] times in the HiFi reads, i.e., at most *t* = 3 standard deviations away from the average coverage. Of those, 11,058,290 are truly uni*k*mers which correspond to a precision of 99.29%; only 79,047 are false positives (0.71% of the total). For *t* = 3, this method recalls 97.84% of the uni*k*mers in the genome. Almost identical results can be obtained from *k* = 17 or *k* = 25. This analysis indicates that selecting *k*-mers that have a number of occurrences in the interval *μ* ± 3*σ* in HiFi reads can recover almost 98% of the true single-copy *k*-mers in the genome with a false positive rate less than 1%. The same analyses carried out on Oxford Nanopore (ONT) reads show that precision, recall, and F1-score for ONT reads are slightly lower than those obtained from PacBio HiFi reads, likely due to the higher rate of sequencing errors in ONT reads (see Figure 10).

Based on this analysis, we used *k* = 21 and *t* = 3 for all the experiments below.

### 2.4 Algorithm

The algorithm used in RAmbler is illustrated in Figure 4. It comprises of six major steps, the first two of which are data preprocessing.

**Figure 4:**
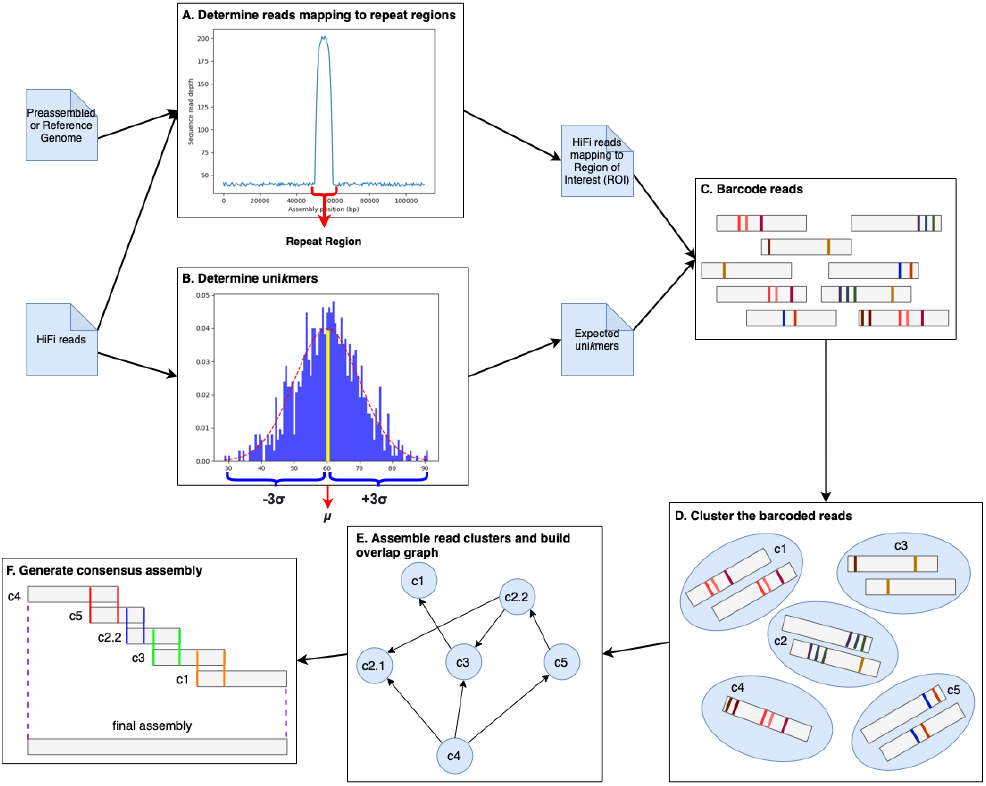
The algorithmic pipeline used in RAmbler

#### A. Determine the reads corresponding to repetitive regions

As mentioned in Section 2.2, RAmbler identifies repetitive regions by mapping all HiFi reads *T* against the draft genome. RAmbler generates the plot of the read coverage across the genome using NucFreq [34]. Unresolved repetitive regions produce distinctive peaks in the coverage plot as illustrated in Figure 2 and Figure 15. Then, RAmbler selects the reads that map to the repetitive regions, as well as the reads extending 50 kb upstream and downstream of the peak, as shown in Figure 4, step A. We call the set of HiFi reads extracted in this step *T*_*r*_, where *r* identifies the repetitive region.

#### B. Determine unikmers

RAmbler uses Jellyfish on the entire set of HiFi reads *T* to obtain the count distribution of 21-mers genome-wide. RAmbler calculates the mean *μ* and standard deviation *σ* of the distribution. RAmbler then selects the 21-mers that fall within the interval [*μ*− 3*σ, μ* + 3*σ*] . According to our analysis in Section 2.3, these 21-mers are true uni*k*mers with high probability (Figure 4, step B).

#### C. Barcode reads

RAmbler uses the set of uni*k*mers to barcode the HiFi reads *T*_*r*_ (Figure 4, step C). RAmbler searches for exact occurrences of the uni*k*mers in the reads *T*_*r*_ or their reverse complement. The set of uni*k*mers present in a read and their location is the *barcode* of that read. For each read, RAmbler stores pairs (*u, j*), where *u* is a uni*k*mer and *j* is the location within the read.

#### D. Cluster the barcoded reads

RAmbler compares the barcode of all pairs of reads to identify shared uni*k*mers. This pairwise comparison allows RAmber to determine which reads are overlapping. Two reads are overlapping if they share at least *th* uni*k*mers, and the set of relative distances between the shared uni*k*mers match within a tolerance up to *to* base pairs. Overlaps are stored in the *barcode graph*: each node in the barcode graph represents a read; nodes in the graph are connected by an edge if the corresponding reads are overlapping according to the criteria described above. Once the graph is completed, RAmbler identifies clusters of reads by determining the connected components of the barcode graph (Figure 4, step D).

#### E. Assemble read clusters and build overlap graph

RAmbler carries out individual local assemblies for each set of clustered reads using a standard HiFi assembler (Hifiasm in this case). Each read cluster is assembled in one or more contigs. RAmbler then uses minimap2 [15] to align assembled contigs to each other. Any contig that is fully contained within another larger contig is removed. RAmbler constructs an overlap graph based on the overlap information provided by minimap2: each node in the overlap graph represents a contig; nodes are connected by edges if they have a suffix-prefix overlap (Figure 4, step E). It is worth noting there could be multiple suffix-prefix overlaps between a pair of contigs. RAmbler retains the overlap with the highest percentage of identity as long as the overlap is at least *mo* = 1000 bps. Furthermore, these suffix-prefix overlaps can occur between the positive or negative strands, resulting in three types of edges. Each edge is labeled by a pair (*t, l*), where *t*∈ {+, −, ∗}, and *l* is the length of the suffix-prefix overlap (Figure 5). Given an edge (*u, v*) in the overlap graph, its type *t* is (i) “+” when there is an overlap between contig *u* and contig *v*, (ii) “−” when there is an overlap between the reverse complement of contig *u* and contig *v*, (iii) “∗” when there is an overlap between contig *u* and the reverse complement of contig *v*.

**Figure 5:**
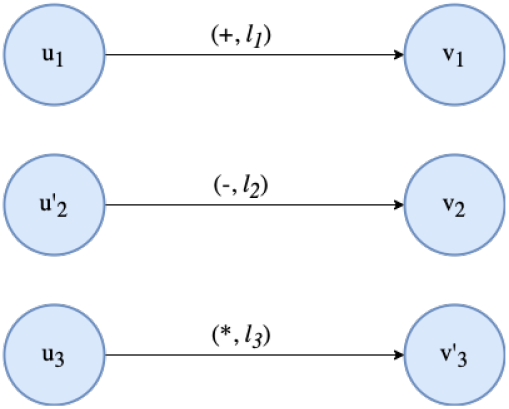
The three types of edges in our overlap graph

#### F. Generate consensus assembly

At this stage, RAmbler needs an estimate of the size of the repetitive region. The estimate can be provided to RAmbler by the user, or it can be obtained from NucFreq plot (Figure 4, step A) as follows. First, RAmbler measures the width of the coverage peak to determine the length of the repeat unit. Then, RAmbler estimates the number of copies by dividing the height of the peak by the average read coverage genome-wide. RAmbler computes the expected length of the repetitive region by multiplying the length of the repeat unit by the number of copies. We add 100 kb to account for the 50 kb upstream and downstream regions.

To compute the final assembly, RAmbler first determines whether the overlap graph is acyclic. If it is acyclic, RAmbler enumerates all possible paths using DFS and generates a set of candidate assemblies. When computing the sequence consensus for suffix-prefix overlaps, if the suffix and the prefix do not match, RAmbler arbitrarily picks the base either from the suffix or the prefix (Figure 4, step F). Among all assemblies, RAmbler selects the one that best matches the estimated length of the repetitive region.

When the overlap graph has cycles, RAmbler partitions the graph into three components: an acyclic pre-cycle subgraph, the cycle itself, and post-cycle subgraph (which could be cyclic), as shown in Figure 6. RAmbler repeats this process iteratively on the post-cycle subgraph until no cycles remain. Once the graph is completely decomposed in a set of acyclic subgraphs, RAmbler generates an assembly for each subgraph as described in the previous paragraph. RAmbler then enumerates all possible combinations of partial assemblies and selects the combination such that the sum of the individual assembly’s length best matches the estimated length of the repetitive region.

**Figure 6:**
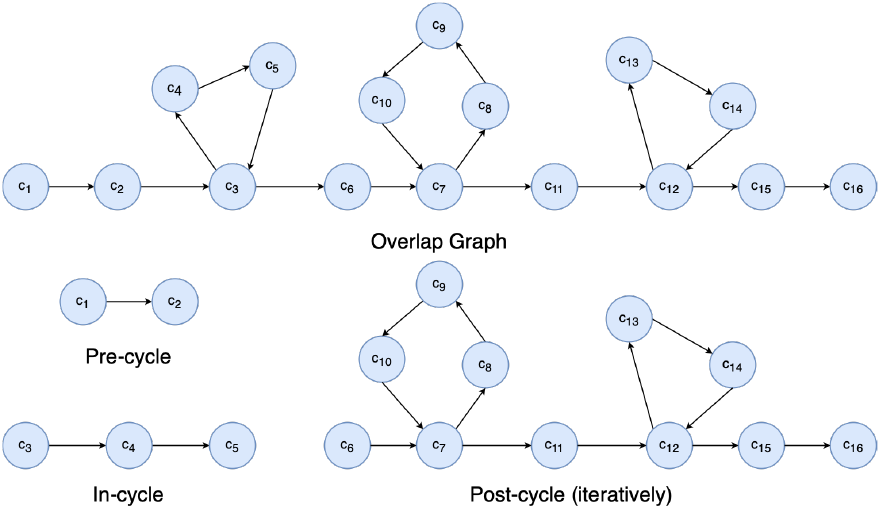
Resolving cycles in an overlap graph

**Figure 7:**
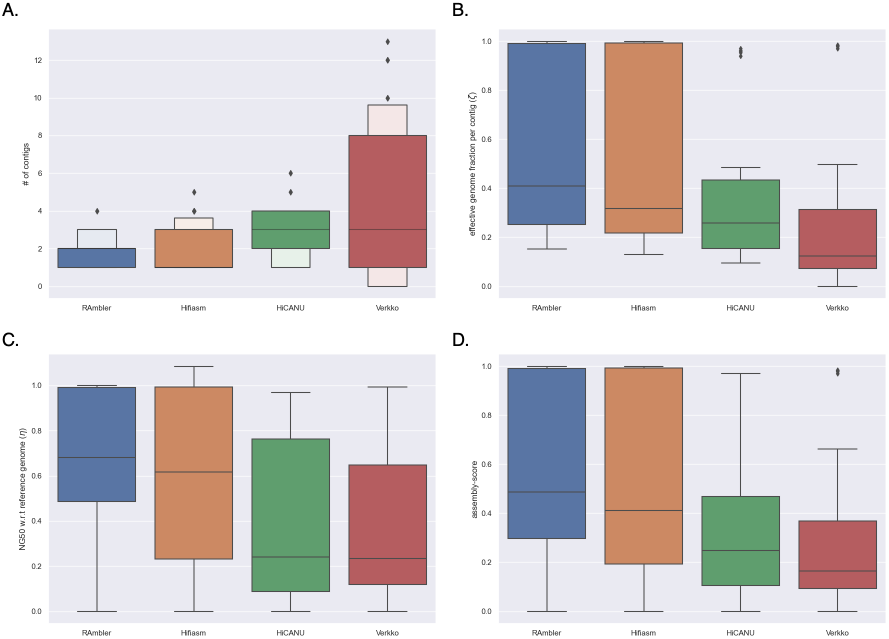
Performance comparison of RAmbler against Hifiasm, HiCANU, and Verkko for 36 different combinations of synthetic data with repetitive regions having repeat sizes = {15, 20} kb, number of copies = {5, 10}, 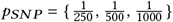 and HiFi reads’ coverage depth = {20, 30, 40}x

**Figure 8:**
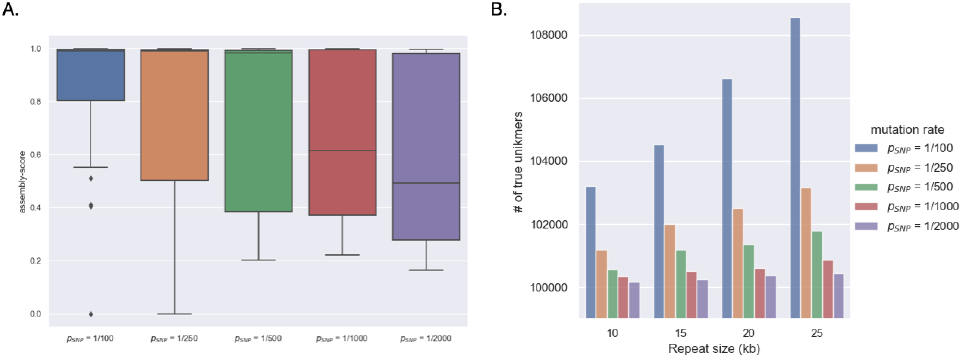
(A) Performance of RAmbler for several choices of the mutation rate *p*_*SN P*_ over 27 different combinations of repetitive regions with repeat sizes = {10, 15, 20} kb, number of copies = {2, 5, 10}, synthetic reads’ coverage depth = {20, 30, 40} x, (B) Number of true uni*k*mers as a function of mutation rate and repeat unit’s sizes (five copies)

A summary of RAmbler’s main parameters *k, t, th, to*, and *mo* with their default values is shown in Table 1.

**Table 1:**
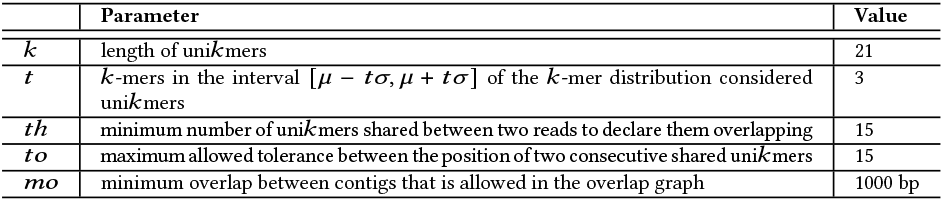
RAmbler’s main parameters.

## 3 EXPERIMENTS AND RESULTS

### 3.1 Synthetic Data Generation

We first generated synthetic repetitive regions using a combination of (i) repeat unit size: 10 kb, 15 kb, 20 kb and 25 kb; (ii) 2, 5, and 10 copies of the repeat unit; (iii) mutation rate in each copy of the repeat 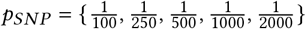. For each combination, we generated HiFi reads using PBSIM on the CCS model with read coverage of 10x, 20x, 30x, and 40x [23]. PBSim requires other parameter values to be set before generating reads, which are provided in Table 2.

**Table 2:** PBSim’s parameters and their values.

| Parameter | Value | Parameter | Value |
| --- | --- | --- | --- |
| Mean read length | 13,500 | Mean read accuracy | 0.998 |
| Standard deviation of read length | 1,500 | Standard deviation of read accuracy | 0.01 |
| Minimum read length | 7,500 | Minimum read accuracy | 0.95 |
| Maximum read length | 20,000 | Maximum read accuracy | 1.0 |

### 3.2 Performance Metrics

Metrics such as NG50, genome fraction, number of contigs, and number of mis-assemblies have been traditionally employed to evaluate the quality of an assembly on synthetic data when the reference genome is known. However, each of these metrics alone does not fully capture all the desired qualities of a “good assembly”. To address this shortcoming we introduce here a new metric called the *assembly score* that summarizes in one number the quality of an assembly in terms of accuracy, contiguity, and completeness. The assembly score is based on two preliminary metrics, as explained next.

#### 3.2.1 Effective genome fraction per contig (ζ)

Consider an assembly that consists of a single contig but contains one mis-assembly. To correct the mis-assembly, the contig needs to be broken, resulting in the creation of an additional contig. Based on this observation we define the *effective number of contigs* as the sum of the number of contigs in the assembly and the number of mis-assemblies. We propose to calculate the effective genome fraction per contig as follows

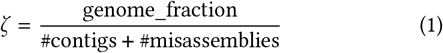

As said, metric ζ takes into account the number of mis-assemblies and penalizes the score accordingly. An assembly that covers 100% of the reference genome (without any mis-assemblies) would yield ζ = 1.0. By considering the effective genome fraction per contig, we can assess the assembly quality while accounting for the presence of mis-assemblies, thereby providing a more comprehensive evaluation.

#### 3.2.2 Normalized NG50 (*η*)

While NG50 is an essential metric for evaluating the contiguity of an assembly, it depends on the size of the reference genome, making it challenging to use it to compare an assembler’s performance across genomes of different lengths. To address this limitation, we normalize NG50 by the size of the reference genome, yielding a metric called *η* defined as follows

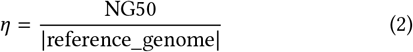

By normalizing NG50 with respect to the reference genome size, *η* is constrained within the range of [0, 1], with a perfect assembly achieving *η* = 1. Observe that it is possible that *η* may exceed 1 when the assembly is over-inflated, i.e., longer than the actual genome. In general, a higher value of *η* indicates a better assembly quality, as long as it is smaller than 1.

#### 3.2.3 Assembly score

The assembly score is defined by computing the harmonic mean of ζ and *η*, as follows

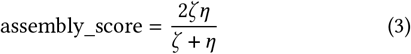

Observe that while ζ is always within 0, 1, *η* can exceed 1, which can result in an assembly score greater than 1. Nevertheless an assembly score closer to 1 indicates a higher quality assembly. This score enables a holistic assessment of the assembly’s quality, taking into account accuracy, contiguity, and completeness, which are all equally important.

### 3.3 Parameter Tuning

As show in Table 1, RAmbler has five parameters. Based on the analysis in Section 2.2, we determined that *k* = 21 and *t* = 3. To find the optimal values for *to* and *th*, we conducted a grid search where *to* = {1, 5, 10, 15, 20 }and *th* = {5, 10, 15, …, 50} (50 combinations). Parameter *mo* (minimum overlap) was set to 1, 000 bps.

RAmbler was tested on 135 synthetic data sets, obtained from the combinations of different choices of the repeat unit size {10, 15, 20} kb, repeat copies {2, 5, 10}, mutation rate 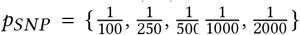, and HiFi reads’ coverage depth 20*x*, 30*x*, 40*x* . Figure 11 shows the assessment results for different metrics (namely number of contigs, number of mis-assemblies, effective genome fraction per contig and normalized NG50), with the best choices for *to* and *th* highlighted in colored rectangles.

When considering the number of contigs (Figure 11-A), observe that RAmbler achieved the best performance for *to* = {15, 20} and *th* = {10, 15}. Regarding the number of mis-assemblies produced by RAmbler (Figure 11-B), the best values were *to* = {15, 20} and *th* = {10, 15, 20, 25 }. In terms of effective genome fraction per contig (ζ) (Figure 11-C), RAmbler had better results with *to* = {10, 15, 20} and *th* ={10, 15, 20 }. Finally, considering the normalized NG50 (*η*) (Figure 11-D), the best outcomes were obtained with *to* = {15, 20} and *th* = 15. By combining all these metrics in the assembly score (Figure 12), we determined that the optimal values are *to* = 15 and *th* = 15.

### 3.4 Experimental Results on Synthetic Data

#### 3.4.1 Comparing RAmbler against Hifiasm, HiCANU, and Verkko

We conducted an extensive performance comparison of RAmbler with other state-of-the-art HiFi assemblers, namely Hifiasm, Hi-CANU, and Verkko, using 36 synthetic data sets. As mentioned in the introduction, SDA is no longer maintained, thus was excluded from this comparison. We generated repetitive regions with two choices of the repeat unit size {15, 20} kb, two choices for the number of copies {5, 10}, and three levels of mutation rate 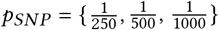 . Synthetic HiFi reads were generated with PBSim with coverage depths of {20*x*, 30*x*, 40*x*} (see Section 3.1 for details).

Figure 7 summarizes the results in terms of four different performance metrics. In Figure 7-A, we compare the number of contigs produced by the different assemblers, where a single contig would be the ideal assembly. Observe that RAmbler consistently produced the lowest number of contigs among all the assemblers. Figure 7-B, C, and D show the results in terms of effective genome fraction per contig (ζ), normalized NG50 (*η*), and assembly score, respectively. For all these metrics, the best assembly is the one that gets closer to 1.0. Observe again that RAmbler achieved higher values on all these metrics compared to the other three assemblers.

#### 3.4.2 Comparing RAmbler against Hifiasm, HiCANU, and Verkko on variable-length repeats

To evaluate RAmbler’s ability to resolve tandem repeats in the presence of variable-length repeat units, we created synthetic repetitive regions in which each repeat copy can vary up to ±5% of the length of a repeat unit. We used five copies of repeat units of {15, 20, 25} kb with mutation rates 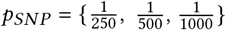. Synthetic HiFi reads with a coverage depth of 30 *x* were generated with PBSim. Table 3 summarizes assembly score results for these nine data sets. RAmbler outperformed other assemblers in 7 out of 9 test runs.

**Table 3:**
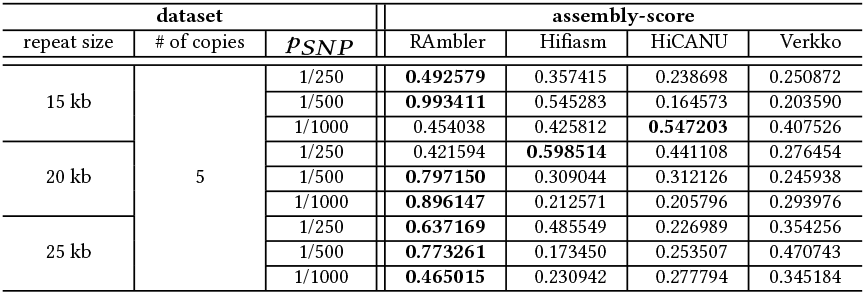
Assembly-score results for nine different data sets of synthetic HiFi reads with coverage 30x, based on a repetitive region with a variable-length (up to ±5% per copy) repeat unit of length {15, 20, 25} kb, and 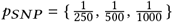; numbers in bolt indicate the best assembly score on each row

### 3.5 Experimental Results on Real Data

Finally, we tested RAmbler on real data for *Arabidopsis thaliana*, using the TAIR10.1 reference genome and PacBio HiFi reads (SRA accession ERR6210723) with an average coverage of 50x. RAmbler detected repetitive regions in chromosomes 2, 3, 4, and 5, which are well-known [3, 8, 20, 36].

In this study, we focused on three centromeric repetitive regions in chromosome 5. The three NucFreq plots in Figure 15 illustrate the mapping coverage in these repetitive regions. RAmbler estimated the first tandem repeat to be composed of 45 copies of a 4.5 kb unit, the second to be 4 copies of a 11.35 kb unit, and the third to be 75 copies of a 10.45 kb unit (Table 4, columns 2 and 3).

**Table 4:** Assembly results obtained by RAmbler, Hifiasm and Verkko on real HiFi reads for three known repeats of *A. thaliana* in chromosome 5; HiCANU failed to complete all these assemblies.

| Repetitive region | Expected |  | RAMbler |  | Hifiasm |  | Verkko |  |
| --- | --- | --- | --- | --- | --- | --- | --- | --- |
|  | unit size | copies | contigs | copies | contigs | copies | contigs | copies |
| chr5:11165kb-11210kb | 4.5 kb | 45 | 5 | 44 | 13 | 0 | 3 | 0 |
| chr5:11300kb-11350kb | 11.35 kb | 4 | 2 | 4 | 7 | 0 | N/A | N/A |
| chr5:11690kb-11720kb | 10.45 kb | 75 | 10 | 72 | 60 | 0 | 82 | 0 |

Reads mapping to these three repetitive regions (including 50 kb upstream and downstream) were given as input to RAmbler, Hifiasm, Verkko and HiCANU. Table 4 shows the number of contigs and the number of copies reconstructed by RAmbler, Hifiasm, and Verkko. We stopped HiCANU after 24 hours of execution on these three inputs, thus it is excluded from the table.

On the first 45-copy tandem repeat, RAmbler generated an assembly with 5 contigs, while Hifiasm created 13 contigs and Verkko produced 3. We aligned these assemblies against the repeat unit extracted from the reference genome using minimap2. The dotplot confirmed 44 copies for RAmbler (dotplot not shown). The assembly produced by Hifiasm and Verkko did not align well against the repeat unit thus resolving zero copies (dotplots not shown).

On the second centromeric 4-copy tandem repeat, RAmbler and Hifiasm generated assemblies with 2 and 7 contigs, respectively. Verkko did not produce any output. The evaluation with minimap2 and dot plot revealed that RAmbler resolved all 4 copies, while Hifiasm failed to resolve any copies of the repeat (dotplots not shown).

On the third, and most challenging, 75-copy tandem repeat, RAmbler, Hifiasm, and Verkko generated assemblies with 10, 60, and 82 contigs, respectively. Among those, RAmbler produced two contigs longer than 100 kb, Hifiasm generated two long contigs, and Verkko produced seven long contigs. When we aligned these long contigs against the repeat unit using minimap2, the dot plot revealed that RAmbler successfully resolved 72 copies, while both Hifiasm and Verkko failed to resolve any copy (dotplots not shown).

Since the “ground truth” is not available for these three repetitive units, we cannot verify whether RAmbler resolved these repeats correctly, i.e., by placing the repeat units in the exact order in which they are supposed to appear. However, our extensive experiments on synthetic data sets have shown that RAmbler is capable of resolving complex tandem repeats with high accuracy.

## 4 CONCLUSION

We presented RAmbler, a novel algorithmic approach for *de novo* genome assembly, specifically aimed at resolving complex repetitive regions. Our method leverages the concept of uni*k*mers, which are single copy *k*-mers, to barcode the HiFi reads. By taking advantage of shared uni*k*mer, RAmbler can reliably select overlaps that would be difficult to be identified by traditional assemblers due to the large number of spurious overlaps in these regions. We also showed the first statistical analysis for the method to identify single copy *k-*mers, and provided a novel comprehensive measure of assembly quality.

Our extensive experiments on more than 140 synthetic data sets and on real HiFi data on three repetitive regions in *A. thaliana*, clearly demonstrate that RAmbler outperforms Hifiasm, Verkko, and HiCANU on reconstructing these repetitive regions.

RAmbler still has some limitations. Its performance drops significantly when 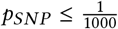(see Figure 8-A). In this case, the copies are almost identical, thus there are not enough uni*k*mers to resolve them. This observation is supported by Figure 8-B, which illustrates that the number of uni*k*mers decreases with *p*_*SN P*_ . However, the mutation rate in real genomes is generally higher than one SNP over a thousand base pairs: in the human genome, for instance, there are at least 10 million SNPs occurring every 100-300 bps [14, 24, 30].

While these initial results are very promising, to establish RAmbler’s true ability in resolving complex repeats, it will have to be tested more extensively on real data for other organisms.

## ACKNOWLEDGMENTS

This project was supported in part by NSF IIS #1814359, NSF CBET #2225878, and NIH 1-R01-AI169543-01 to SL.

## A SUPPLEMENTAL FIGURES AND TABLES

**Figure 9:**
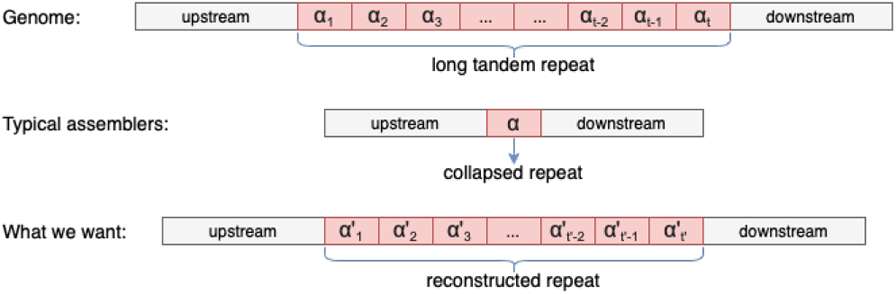
The problem formulation: tandem repeats *α*_1_, *α*_2_, …, *α*_*t*_ are over-collapsed by traditional assemblers; we want to reconstruct all the copies of the repetitive regions

**Figure 10:**
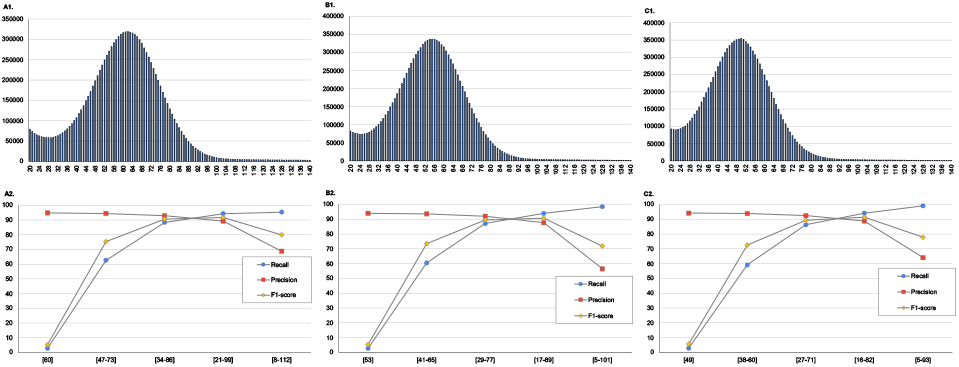
Oxford Nanopore *k*-mer distribution for *k* = 17 (A1), *k* = 21 (B1) and *k* = 25 (C1) for *Saccharomyces cerevisiae* dataset (SRA accession SRR 15597407); precision, recall, and F1-score for uni*k*mers when *k* = 17 (A2), *k* = 21 (B2) and *k* = 25 (C2) for larger and larger intervals centered at the average sequencing coverage

**Figure 11:**
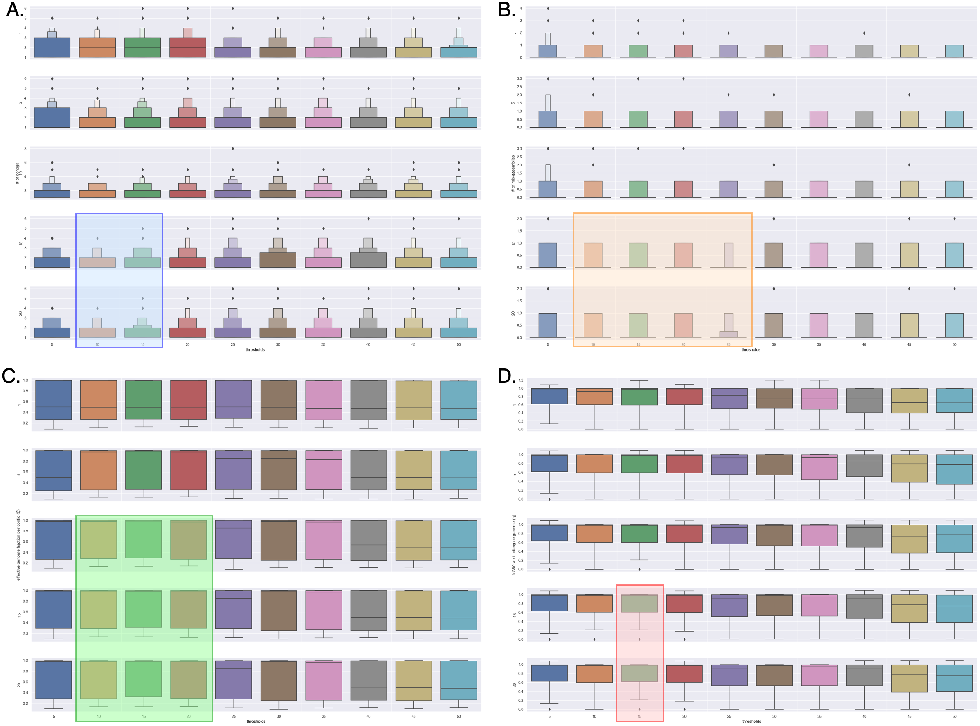
Performance of RAmbler for difference combinations of tolerance *to* = {1, 5, 10, 15, 20}, threshold, *th* = {5, 10, 15, 20, 25, 30, 35, 40, 45, 50}, repeat size = {10, 15, 20} kb, number of copies = {2, 5, 10}, 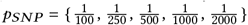, on synthetic reads with coverage depth = {20, 30, 40} x; the rows correspond to different values of columns correspond to different values of *th*; (A): number of contigs, (B): number of mis-assemblies, (C): effective genome fraction per contig ζ, and (D): normalized NG50 *η*.

**Figure 12:**
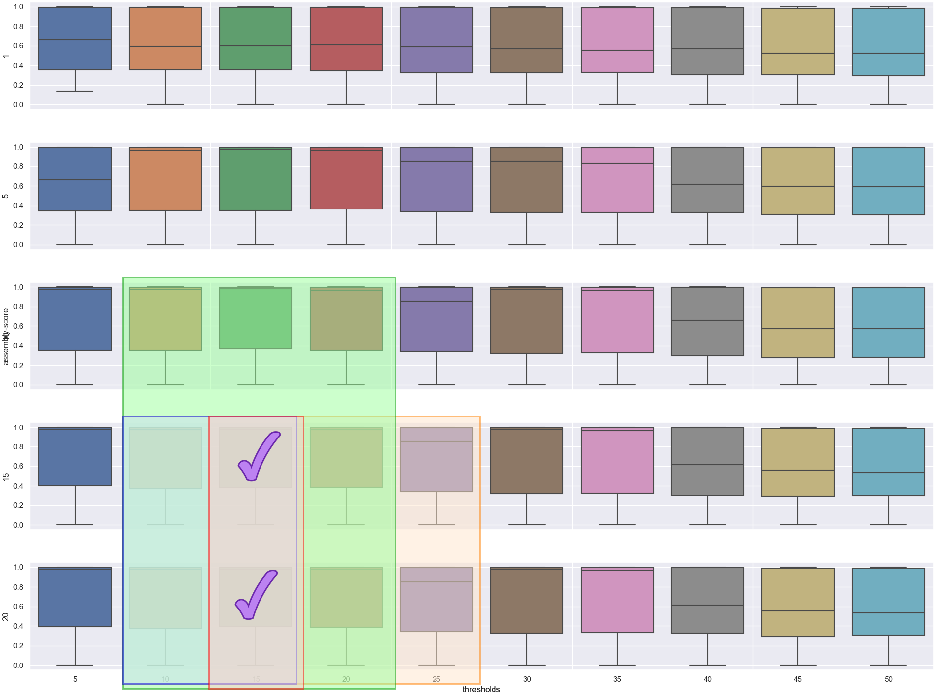
Assembly scores achieved by RAmbler for different combinations of tolerance *to* = {1, 5, 10, 15, 20}, threshold *th* = {5, 10, 15, 20, 25, 30, 35, 40, 45, 50}, repeat size = {10, 15, 20} kb, number of copies = {2, 5, 10}, 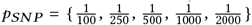, on synthetic reads with coverage depth = {20, 30, 40 }x. The rows correspond to different values of *to*, the columns correspond to different values of *th*.

**Table 5:**
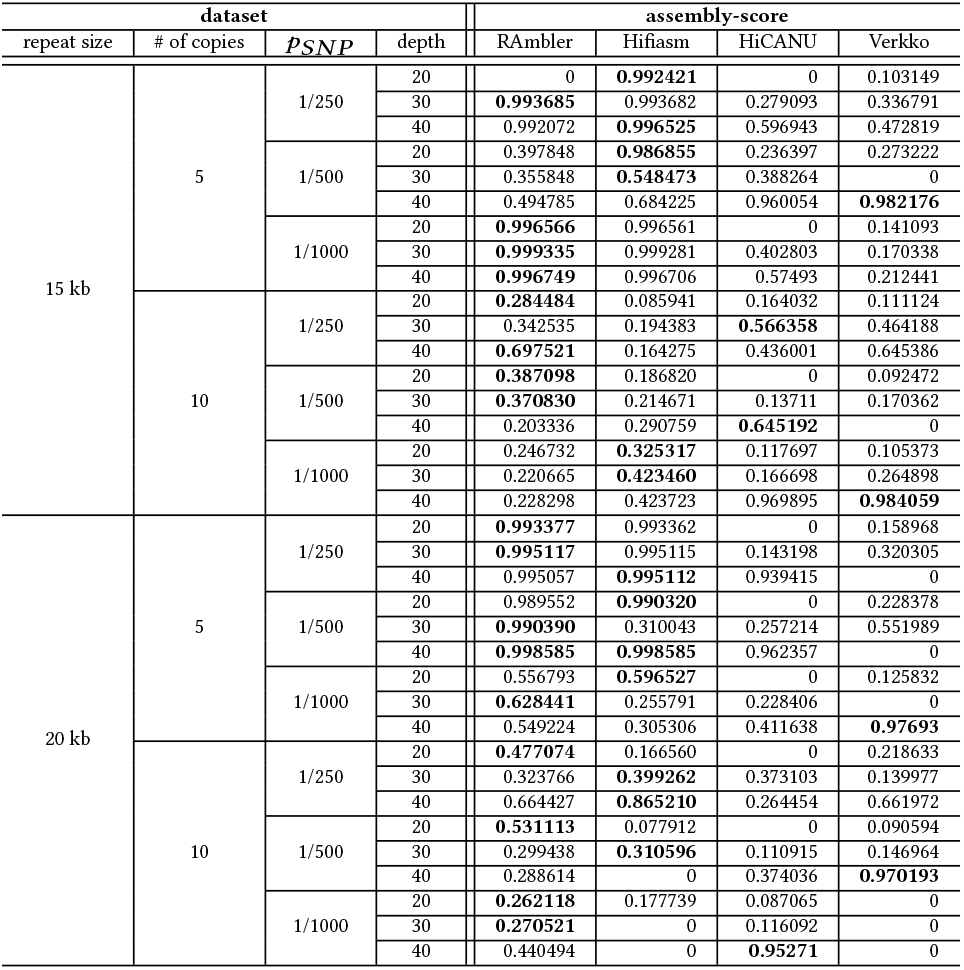
Assembly-score of RAmbler, Hifiasm, HiCANU, and Verkko on 36 synthetic data sets; numbers in bolt indicate the best assembly score on each row.

**Figure 13:**
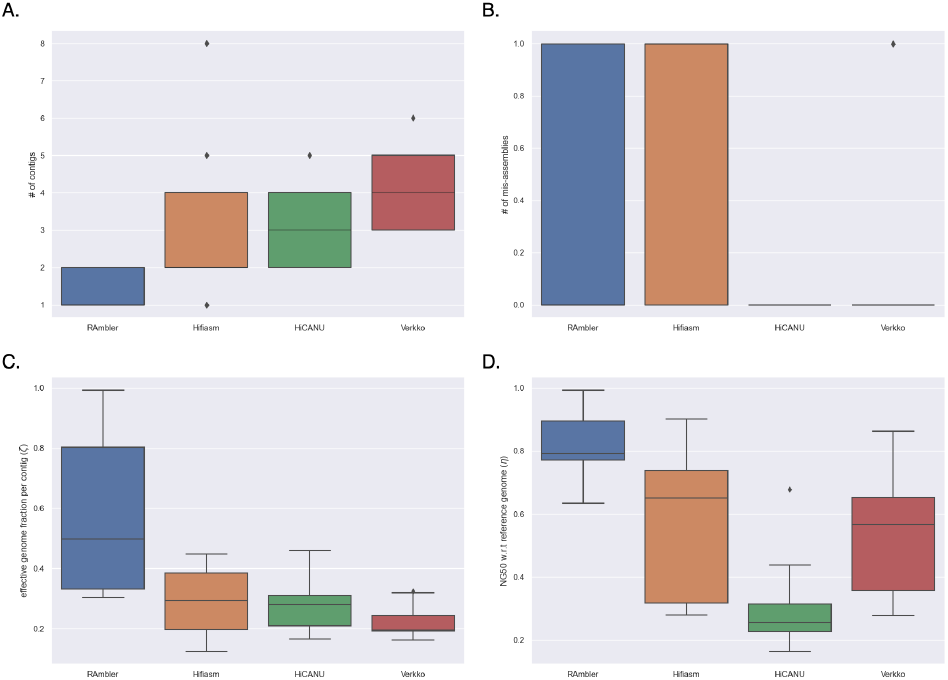
Performance of RAmbler, Hifiasm, HiCANU and Verkko on synthetic data with 30x coverage depth for nine different combinations of repetitive regions containing five copies of a repeat size = {15, 20, 25} kb and variable-length (up to ±5% per copy), 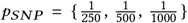: number of contigs, (B): number of mis-assemblies, (C): effective genome fraction per contig ζ, and (D): normalized NG50 *η*.

**Figure 14:**
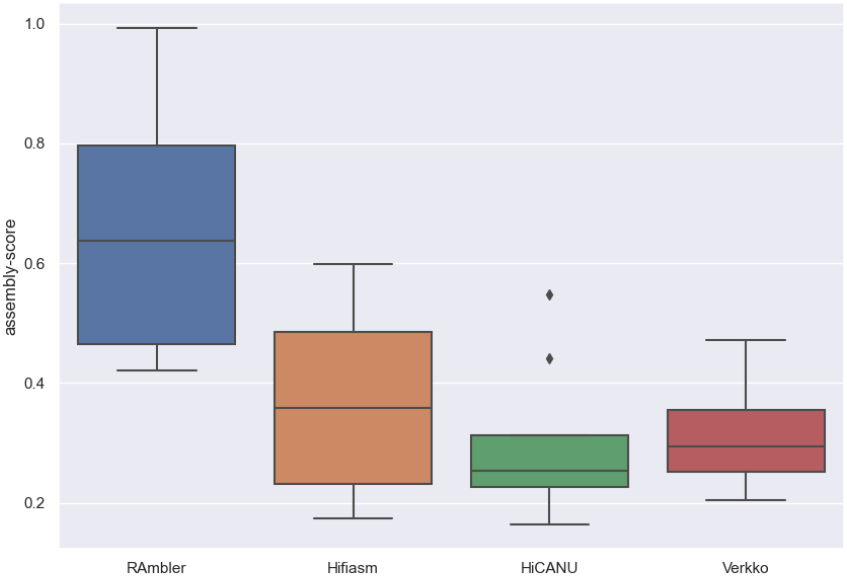
Assembly-score of RAmbler, Hifiasm, HiCANU and Verkko on synthetic data with 30x coverage depth for nine different combinations of repetitive regions containing five copies of a repeat size = {15, 20, 25} kb and variable-length (up to ±5% per copy), 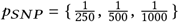.

**Figure 15:**
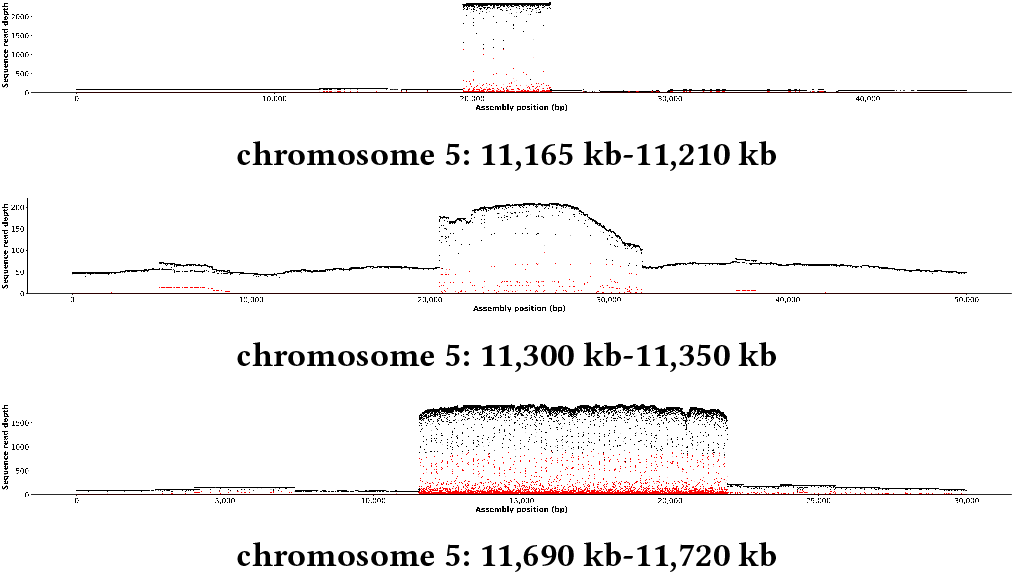
PacBio HiFi mapping coverage for chromosome V in *A. thaliana* illustrated using NucFreq; the coverage spikes indicate the presence of repetitive regions

